# Rat hepatitis E virus infection has multiphasic viral replication kinetics *in vivo*

**DOI:** 10.64898/2026.06.03.728993

**Authors:** Zhenzhen Shi, Xin Zhang, Niels Cremers, Johan Neyts, Harel Dahari, Suzanne J. F. Kaptein

## Abstract

**Background and Aims:** Hepatitis E virus (HEV) infections are a growing threat to global public health. To obtain an in-depth understanding of HEV infections in untreated and ribavirin-treated rats, we characterized the early HEV viral kinetics using rat HEV (rHEV) as a surrogate model and using two routes of virus inoculation: intravenous (I.V.) or oral infection.

**Approach and Results:** We frequently collected feces, serum, and tissue samples up to 60 days after infection in both infection models to characterize the rHEV viral kinetics. A ~2-week delay in quantifiable RNA levels in feces was observed in the oral versus the I.V. infection model. Early rHEV viral kinetics in feces were found to be multiphasic and showed good concordance with those in the various tissue compartments studied. Comparison of the viral kinetics in these samples also revealed that the liver may serve as the initial site of rHEV replication, followed by replication in the intestine and spleen. While a dosage of 60 mg/kg/day ribavirin was found optimal to maintain rHEV RNA levels at (nearly) undetectable in feces, levels were detectable in the liver and increased both in feces and liver after treatment discontinuation.

**Conclusions:** We found that the two rHEV infection models share similar multiphasic viral kinetics with the liver as the main site of viral replication. Additionally, the rHEV RNA load in feces could be used as a reliable proxy for that in the liver, spleen, and intestine. We also show that ribavirin at 60 mg/kg/day was partially effective in preventing viral rebound. These findings may aid in exploring the correlation between the infection phases and antiviral efficacy, ultimately guiding therapeutic decisions.

## INTRODUCTION

Hepatitis E virus (HEV) ranks amongst the leading causes of acute viral hepatitis disease. HEV is responsible for approximately 20 million infections, resulting in an estimated 3.3 million symptomatic HEV infections and 70,000 fatalities, annually [1, 2]. Although acute HEV infections usually are self-limited in immunocompetent individuals, immunocompromised patients, including organ transplant recipients or patients with HIV, can develop persistent chronic infections, thereby increasing the risk of rapid progression to fibrosis and cirrhosis [3]. Furthermore, an HEV infection during pregnancy is linked to increased risks, such as elevated incidences of premature delivery, stillbirths, and maternal mortality [4].

The *Paslahepevirus* genus, which belongs to the subfamily *Orthohepevirinae* (family *Hepeviridae*), is commonly associated with human HEV infections [5]. Within this genus, genotype (GT) 1, 2, 3 and 4 are most frequently linked to human HEV infections. GT1 and 2 are associated with human infections only and are transmitted fecal-orally via contaminated drinking water, whereas GT3 and 4 are zoonotic, and are transmitted through contaminated food and blood products [6, 7]. Since the first case of transfusion-transmitted HEV (TT-HEV) infection reported in 2004 [8], there have been successive reports in Japan [9] and Europe [10-12]. Although most TT-HEV infections remain asymptomatic, they can lead to severe clinical outcomes, particularly in immunocompromised individuals and pregnant women [13, 14].

Human cases of rat HEV (*Rochahepevirus ratti*) infections are increasingly reported worldwide [15-18]. Similar to human HEV infections, patients with underlying disease(s) or organ transplant recipients are at risk of developing serious liver disease when infected with rat HEV (rHEV). Transmission of rHEV to humans via the fecal-oral route [19] might occur through direct infection or via alternative exposure pathways, which include contact with environmental surfaces contaminated by rat droppings [20] or transmission through intermediate hosts such as pigs, cats, and dogs [21, 22]. However, these transmission modes require further verification.

Ribavirin (RBV) is widely used off-label for treating both human and rat HEV infections [18], albeit with limited effectiveness and significant adverse effects [23]. Alternative treatment options also come with their limitations and drawbacks, e.g., only applicable in a restricted patient population, not efficacious as monotherapy, the emergence of drug-associated mutations [24]. Therefore, there is a great need to better understand the viral kinetics of HEV through different transmission routes, which could aid in assessing the efficacy of novel/repurposed antiviral drugs. We previously implemented two rHEV infection models using athymic nude rats in which rats were inoculated via different routes: (I) intravenous (I.V.) inoculation to mimic TT-HEV infection [25], and (II) oral inoculation with an infectious feces suspension to mimic fecal-oral transmission [19]. However, a detailed analysis of the rHEV infection kinetics in these rat infection models is currently lacking. Here, we aim to explore the viral kinetics of rHEV in both rHEV infection models, and to elucidate the viral dissemination following I.V./oral infection. Frequently collected fecal and tissue samples enabled us to characterize rHEV viral kinetics in feces and investigate its correlation with viral kinetics in various tissue compartments. We also evaluated the effectiveness of a 12-day RBV treatment regimen in suppressing rHEV replication in both infection models to understand the link between the rHEV viral kinetics and RBV treatment success/failure.

## METHODS

### Compounds

Ribavirin was purchased as Virazole from Valeant Pharmaceuticals and dissolved freshly in PBS at 15, 30, or 45 mg/mL every day for use in the *in vivo* experiments.

### Animal experiments

Four- to six-week-old homozygous female athymic nude Hsd:RH-*Foxn1*^rnu^ rats (*Rattus norvegicus*; Inotiv, Horst, The Netherlands) were used in the animal experiments. Housing conditions and experimental procedures were approved by the KU Leuven ethics committee (P003/2021), following institutional guidelines approved by the Federation of European Laboratory Animal Science Associations (FELASA). Rats were housed per two in individually ventilated cages (cage type GR900, Sealsafe Plus, Tecniplast) at 21 °C, 55% humidity and 12:12 light:dark cycles. Rats were provided with food and water *ad libitum*, a cardboard house, and wooden gnawing blocks.

### Viral kinetics experiments in untreated rats

#### Oral infection model

Athymic nude rats (*n*=20) received oral administrations of a diluted infectious liver homogenate (500 μL per rat, comprising approximately 2×10^7^ viral RNA copies) on day 0 and day 2 p.i.. To track the viral kinetics of rat HEV in various organs at different time points after oral infection, feces and serum were collected every 5 or 10 days for viral RNA quantification, respectively. Rats (*n*=4) were euthanized via i.p. injection of dolethal on day 5, 10, 20, and 30 p.i,. On day 60 p.i., only 1 rat was humanely killed while the other three rats in this group were included in an experiment that was not part of this study. As a consequence, serum and feces were collected from all 4 rats on day 60 p.i., whereas intestine and spleen samples were collected from 1 rat. Viral RNA levels in feces, serum, liver (two pieces) and other tissues (intestine, spleen) were quantified by means of RT-qPCR, as described previously [19].

#### I.V. infection model

Athymic nude rats (*n*=10) were injected intravenously in the tail vein with 200 µL of a diluted liver homogenate of rat HEV strain LA-B350 (corresponding to approximately 2×10^7^ viral RNA copies), as earlier described [25]. Ten rats were infected to monitor the kinetics of viral replication in feces. Fecal samples were collected every 2-3 days until day 26 p.i. for viral RNA detection via RT-qPCR.

### Viral kinetics experiments in rats treated with ribavirin

#### Oral infection model

Athymic nude rats (*n*=16) received oral administrations of a diluted infectious feces suspension (500 μL per rat, comprising approximately 2×10^7^ viral RNA copies) on day 0 and day 2 p.i.. Rats were orally administered ribavirin (60 mg/kg) once daily for 12 days, starting 2 hours before the infection on day 0 and continuing until day 11 p.i.. Rats were weighed and checked for clinical signs throughout the experiment. Feces samples were collected on various time points p.i.. Rats were humanely killed on day 30 p.i. (i.e., 18 days after discontinuation of ribavirin treatment; *n*=8 per group) after collecting the last stool samples.

#### I.V. infection model

To assess the efficacy of different doses of RBV in the I.V. infection model (see above for the infection procedure in the I.V. infection model), athymic nude rats (*n*=20; *n*=5 per group) were administered orally with either a control vehicle (PBS) or RBV at doses of 30, 60, and 90 mg/kg, 2 hours prior to infection and treatment continued once daily until day 10 p.i. (RBV 90 mg/kg) or day 11 p.i. (vehicle and RBV at 30 or 60 mg/kg). Throughout the experiment, rats were weighed and checked for clinical signs every day. Fecal samples were collected every 4 days for quantification of the viral RNA load by RT-qPCR. On day12 p.i., rats were euthanized and livers were harvested and stored at –80□°C until further use (viral RNA quantification via RT-qPCR).

### Characterization of HEV RNA kinetics in feces

The HEV RNA kinetics were categorized into distinct groups by empirical analysis. Each identified slope was calculated by linear regression as previously [26]. We defined a LLOQ phase if the viral load is below lower limit of quantification. A plateau phase was defined if the difference between viral loads at two consecutive timepoints is less than 0.5 log_10_ or when the slope calculated from three consecutive viral loads is not significantly greater than 0 (*p*-value >0.05 by linear regression). An ascension phase was identified in cases where the slope was significantly greater than 0 (*p*-value <0.05 by linear regression) or the difference in two subsequent viral loads is greater than or equal to 0.5 log_10_ RNA copies.

### Statistical analysis

Statistical analysis was performed with R software (R version 4.3.2). *lm* function was used to calculate the slope of linear regression model which fitted to the experimental data. *p*-values <0.05 indicate that the slopes are significantly different than 0. The two-sample Mann-Whitney *U* test was employed to evaluate differences in two groups with different sample sizes. Kruskal-Wallis Test was employed to evaluate differences in multiple groups (>2 groups) with different sample sizes. *p*-values <0.05 were regarded as statistically significant difference.

## RESULTS

### rHEV viral kinetics in the feces following oral infection is multiphasic

We first investigated the kinetics of rHEV RNA in the feces in the oral infection model, which simulates fecal-oral transmission. In brief, athymic nude rats (*n*=20) were orally inoculated twice (i.e., on day 0 and 2 post-infection [(p.i.]) with liver homogenate containing approximately 2×10^7^ viral RNA copies, essentially as described previously [19]. Fecal samples and serum were collected for viral RNA detection every 5 days (feces) or 10 days (serum). Additionally, different organs (liver, intestine, and spleen) were harvested on day 5, 10, 20, 30, and 60 p.i. (*n*=4 for each indicated timepoint, except for day 60 p.i. [*n*=1]; Figure 1A).

**FIGURE 1.**
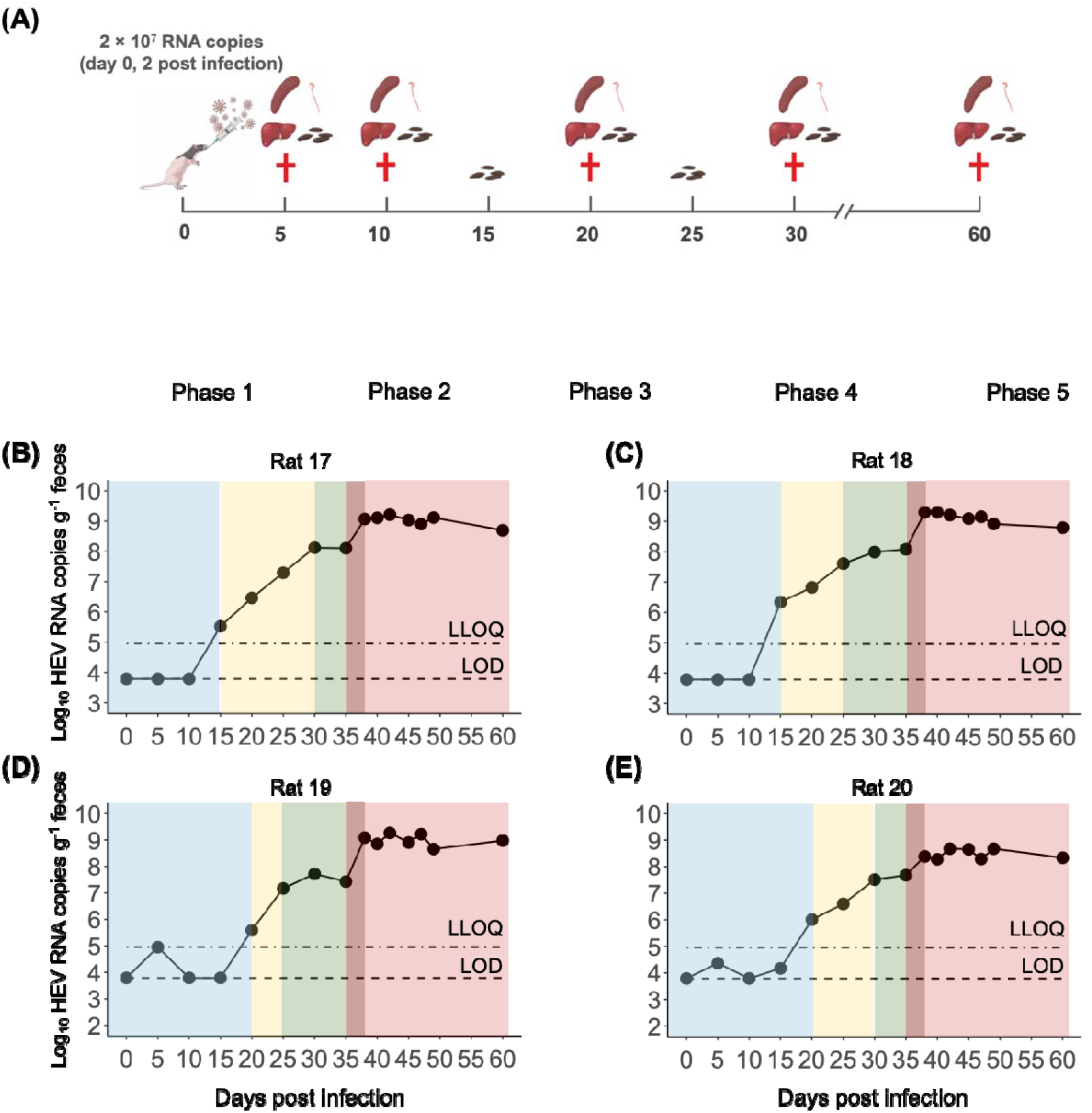
rHEV RNA kinetics in feces of rats in the oral infection model. (A) Schematic illustrating the experimental framework for studying viral kinetics of rHEV using an oral infection model. Rats (*n*=20) were infected orally with rat HEV liver homogenate on day 0 and 2 p.i.. Four rats were euthanized for viral RNA detection in the tissues (i.e., liver, intestine, and spleen) on day 5, 10, 20, 30 p.i., and one rat was euthanized on day 60 p.i. [the other three rats in this group were subsequently included in an experiment that was not part of this study]. (B-E) rHEV RNA kinetic patterns in the feces of four representative rats (all euthanized on day 60 p.i.). Blue shading, yellow shading, green shading, dark red shading, and red shading areas represent LLOQ (Phase 1), rapid ascension (Phase 2), plateau (Phase 3), rapid ascension (Phase 4), and plateau (Phase 5), respectively. The lower limit of quantification (LLOQ) is represented by a dotted-dashed line, and the limit of detection (LOD) is represented by a dashed line.

We identified five phases in the rHEV RNA kinetics in the feces (Supplemental Table S1; Figure 1B-E depicts the viral kinetics in the feces of four representative rats; Supplemental Figure S1 shows the viral kinetics in the feces of all infected rats). In the initial Phase 1 (or lower limit of quantification [LLOQ] phase), viral RNA levels in the feces of infected rats were below the LLOQ, which lasted 17.5 ± 2.9 days (blue shading in Figure 1B-E; Supplemental Table S1). Phase 2 commenced when viral RNA levels in the feces first equaled or surpassed the LLOQ. This phase, also called the ascension phase, was characterized by a rapid increase in HEV RNA levels (yellow shading in Figure 1B-E), as illustrated by an increase in the mean slope by 0.19 ± 0.09 log_10_/day (Supplemental Table S1; *p*-value <0.05) and by an average doubling time (t_2_) of ~4 days (4.0 ± 1.4 days, from day 17.5-27.5 p.i.). In the next phase (Phase 3), the mean slope of viral kinetics reduced to −0.013 ± 0.011 log_10_/day (*p*-value >0.05, indicating a slope not different than zero), indicative of a transition in the viral kinetics from an ascension phase to a plateau phase with an average HEV RNA level of 7.6 ± 0.4 log_10_ copies g^-1^ from day 27.5 to day 35 p.i. (green shading in Figure 1B-E). Phase 3 is followed by another ascension phase (Phase 4; dark red shading in Figure 1B-E) that was characterized by a higher mean slope of 0.38 ± 0.14 log_10_/day with a shorter t_2_ of ~2 days (2.0 ± 0.8 days) from day 35-38 p.i. as compared to Phase 2 (Supplemental Table S1). Finally, the viral kinetics entered a second plateau phase (Phase 5; red shading in Figure 1B-E), which persisted for 22 days (day 38-60 p.i.) and with mean viral RNA levels reaching a plateau that was approximately 1 log_10_ higher (8.7 ± 0.5 log_10_ copies g^-1^ feces; Supplemental Table S1) than in Phase 3.

### rHEV viral kinetics in feces strongly correlates with those in liver, intestine, spleen, and serum

Given the association of HEV infection with a wide spectrum of extrahepatic manifestations, we explored the dissemination of rHEV to various tissues (liver, intestine and spleen) following an oral infection. On day 5 p.i., viral RNA in the examined tissues was undetectable in all rats (Figure 2, Figure 3). On day 10 p.i., two out of four (50%) rats showed viral RNA levels above the LLOQ in the liver (median level 5.3 log_10_ copies g^-1^; min-max: 5.2-5.3 log_10_) while viral RNA levels were undetectable in the intestine and spleen samples from all rats. From day 10 to 30 p.i., the median viral RNA levels in the liver increased from 5.3 log_10_ copies g^-1^ (min-max: 5.2-5.3 log_10_) to 9.8 log_10_ copies g^-1^ (min-max: 9.5-10.0 log_10_). Elevated viral RNA levels were also noted in the intestine and spleen, though they remained lower than in the liver on day 30 p.i.. More specifically, viral RNA in the intestine increased from LLOQ to 7.3 log_10_ copies g^-1^ (min-max: 7.0-7.6 log_10_) and in the spleen from LLOQ to 7.9 log_10_ copies g^-1^ (min-max: 6.9–8.1 log_10_) on day 30 p.i.. Viral RNA levels in the serum remained the lowest compared to those in the studied tissues, with all rats being positive on day 30 p.i. (Figure 3). On day 60 pi, feces and serum were collected from all four rats in this group, while liver, intestine, and spleen samples were collected from only one rat as the other the three rHEV-infected rats were included in a different experiment that was not part of this study (and were thus not euthanized). The viral RNA load in liver, intestine, spleen and serum samples were 9.1 log_10_ copies g^-1^, 7.5 log_10_ copies g^-1^, 7.6 log_10_ copies g^-1^, and 5.9 log_10_ copies ml^-1^, respectively. These levels are comparable to earlier obtained rHEV RNA levels on day 60 p.i. [19], and show that viral RNA levels had reached a steady state in all the studied tissues. When examining the correlation of viral RNA kinetics in the feces, liver, intestine, spleen, and serum, the viral RNA kinetics in the feces strongly correlated (Pearson correlation coefficients of 0.94-0.99) with those in the other studied compartments (liver, intestine, spleen, and serum) from day 20 p.i. onwards (Figure 3A-C, Supplemental Figure S2). Notably, viral RNA levels in the liver were about 10- to 100-fold higher than in the intestine and spleen on almost all studied days, indicating that the liver is the main site of rHEV replication (Figure 2).

**FIGURE 2.**
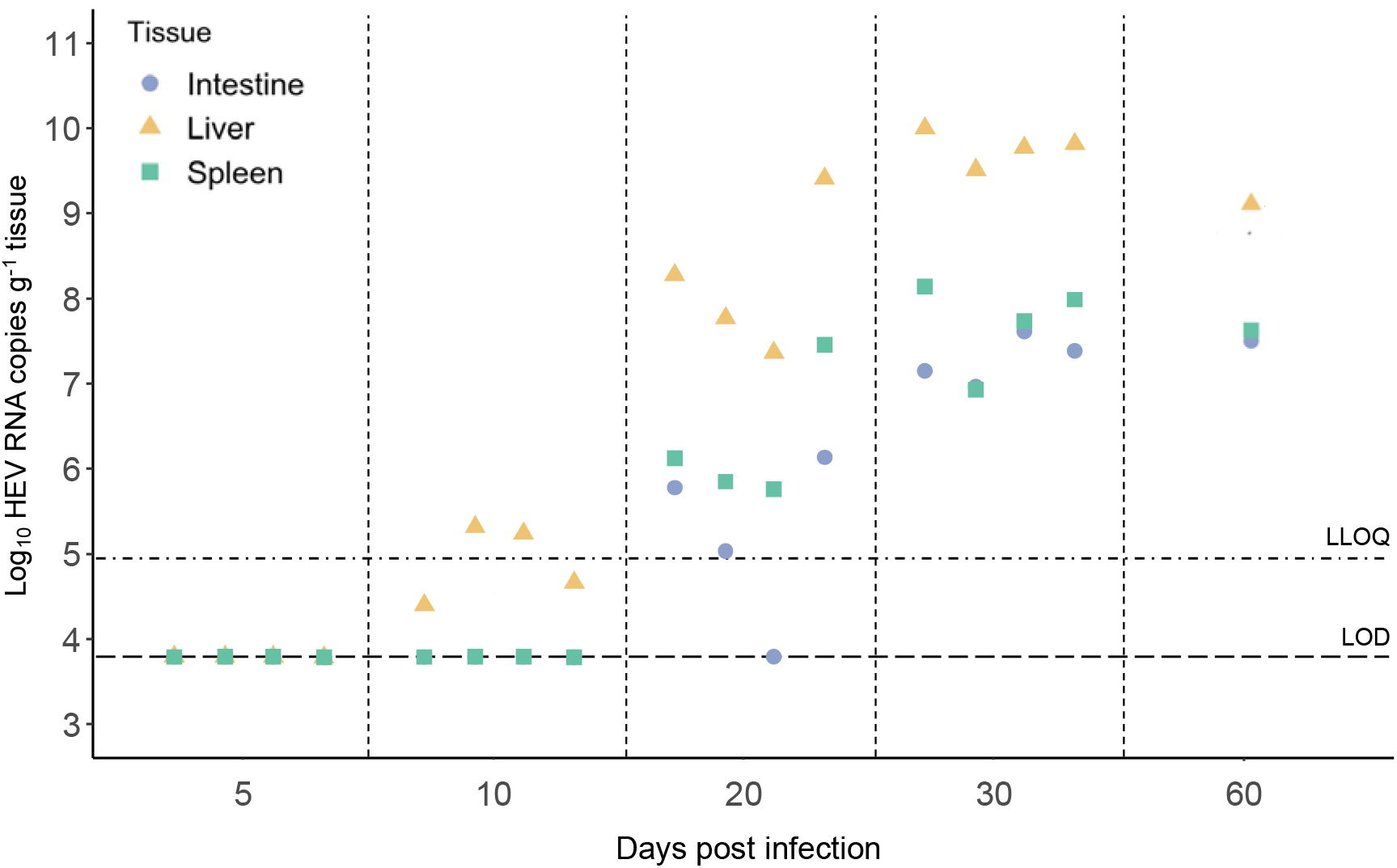
rHEV RNA kinetics in various tissues at different days post oral infection. Solid triangles, circles, and squares that are arranged vertically along the same line represent the liver, intestine, and spleen samples, respectively, collected from the same rat (*n*=17 in total). Four rats were sacrificed to measure the tissue samples on day 5, 10, 20, 30 p.i., and 1 rat was sacrificed on day 60 p.i. [the other three rats in this group were subsequently included in an experiment that was not part of this study]. The lower limit of quantification (LLOQ) and the limit of detection (LOD) for tissues are indicated by different types of dashed lines.

**FIGURE 3.**
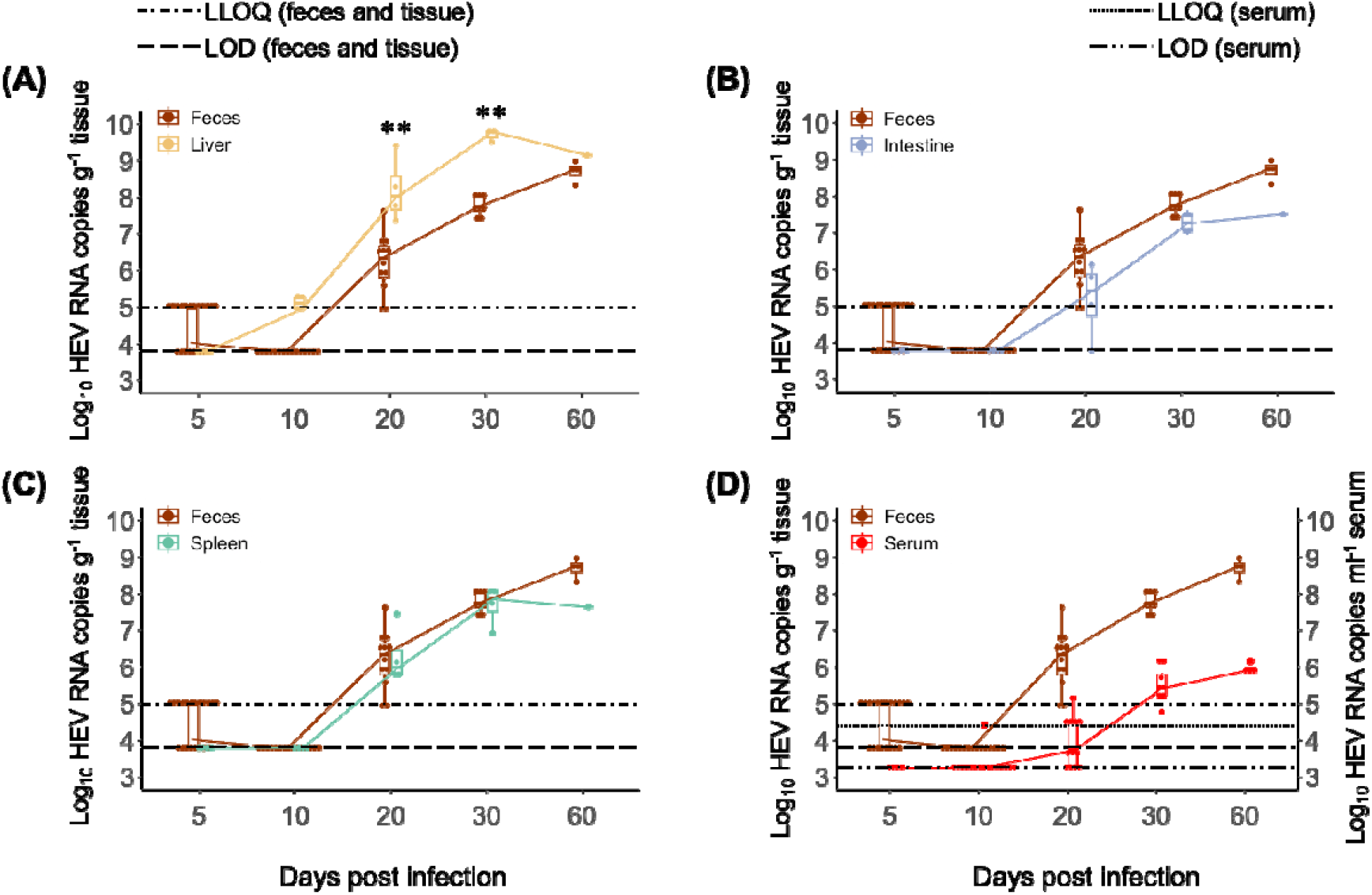
Comparison of rHEV viral kinetics in feces and different tissue samples/serum in the oral infection model. (A-D) The boxplots illustrate the interquartile range (boxed area: 25^th^ to 75^th^ percentiles); the median value of the group is indicated by the central line within the box. Each dot corresponds to the data from an individual rat. A solid line that connects the median values across the boxplots reflects the trajectory of rHEV kinetics. On day 60 pi, feces and serum were collected from all 4 rats belonging to this group while tissue samples (liver, intestine, spleen) were collected from only one rat [the other three rats in this group were included in an experiment that was not part of this study]. The lower limit of quantification (LLOQ) and the limit of detection (LOD) for feces/tissue and serum are indicated by different types of lines. Any values between LOD and LLOQ were set to LLOQ for the purpose of producing boxplots, which was applied to all boxplots. The boxplots thus represent the minimal estimate of the range of the viral RNA levels when there are values below the LLOQ. **, *p* <0.01.

### Similar early rHEV RNA kinetics in the I.V. versus the oral infection model

Although HEV is primarily transmitted through the fecal-oral route, a proportion of the HEV infections occurs through blood transfusion. We therefore also explored the rHEV RNA kinetics in the TT-HEV infection model. In brief, athymic nude rats (*n*=10) were intravenously infected with 1% rat HEV liver homogenate, as previously described [25]. Fecal samples were collected every 1-2 days until day 26 p.i. and processed for viral RNA quantification via RT-qPCR. Interestingly, analysis of viral RNA shed in feces revealed three distinct kinetic patterns similar to those observed in the oral infection model. These patterns included the LLOQ phase, the ascension phase, followed by a plateau phase (Figure 4; Supplemental Table S2). In comparison to the oral infection model, Phase 1 (LLOQ) lasted an average of 4.0 ± 2.2 days, which is 4 times shorter than Phase 1 in the oral infection model. The LLOQ phase was succeeded by Phase 2 (ascension phase) with an average t_2_ of ~4 days (4.1 ± 0.8 days, from day 4.0-17.3 p.i.) and a mean slope of 0.17 ± 0.03 log_10_/day, akin to Phase 2 in the oral infection model. Viral RNA levels in the feces then culminated to a plateau (7.8 ± 0.5 log_10_ copies g^-1^; Supplemental Table S2) in Phase 3, which lasted from day 17.3-25 p.i., akin to Phase 3 in the oral infection model. All in all, the early phase rHEV kinetics (Phase 1-3) in the feces are similar in both infection models.

**FIGURE 4.**
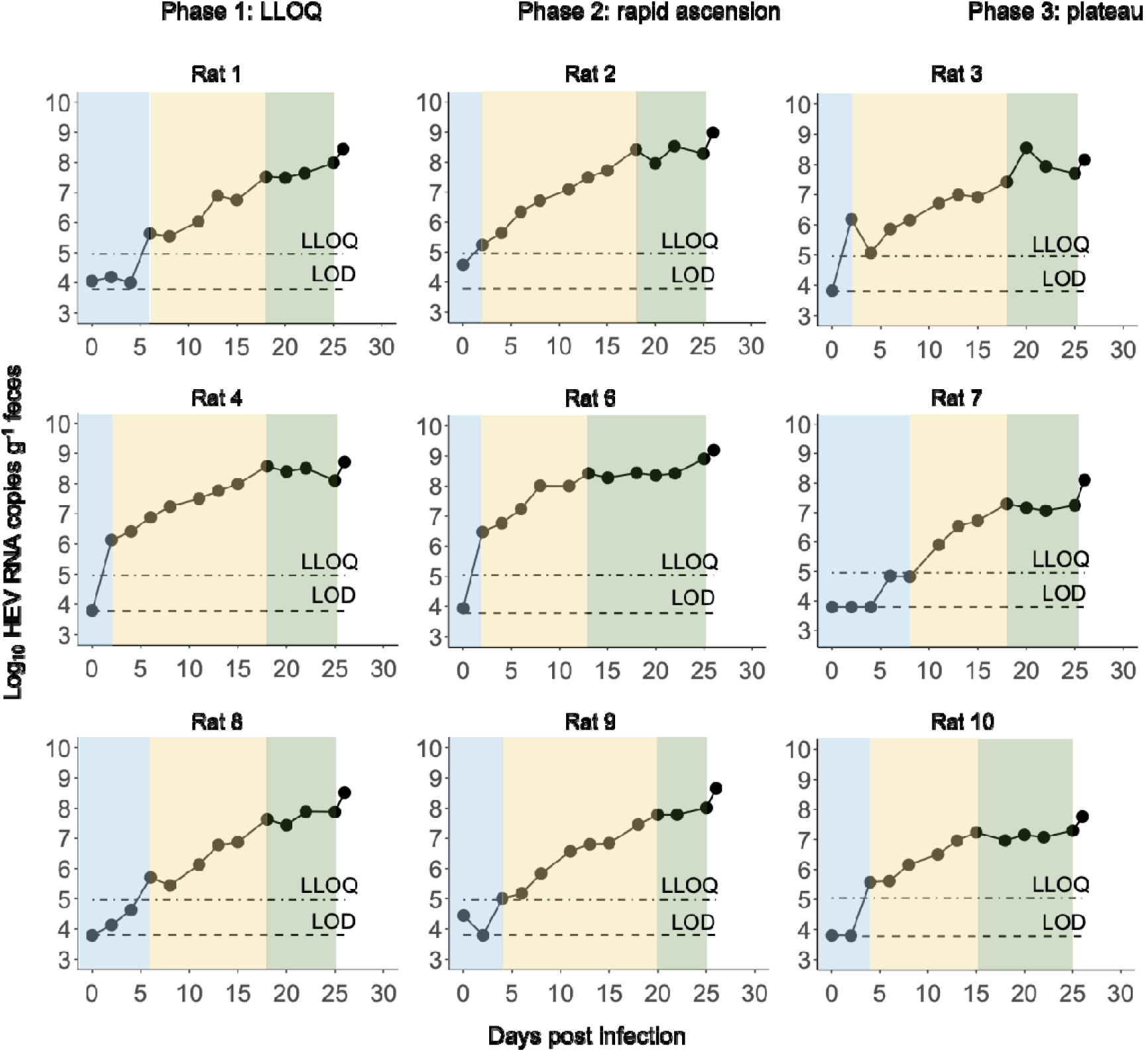
Viral kinetic patterns of rHEV in the feces in the I.V. infection model. Viral RNA levels in the feces at different time points from rats (*n*=10) after I.V. infection with rHEV liver homogenate. Blue shading, yellow shading, and green shading areas represent Phase 1, Phase 2, and Phase 3, respectively. Rat 5 has its distinct kinetic pattern (staircase, as shown in Supplemental Figure S4). The last timepoint (day 26 p.i.) was omitted for rat 1 and rat 6 due to the slope of the linear regression model being significantly different than 0 (*p*-value <0.05) (Supplemental Table S1), suggesting that another ascension phase may arise. LLOQ (dotted-dashed line) and LOD (dashed line) indicate a lower limit of quantification and limit of detection, respectively.

### Assessing the optimal ribavirin dose through rHEV kinetics

We first assessed the effect of a once-daily 60 mg/kg dose of RBV on the rHEV kinetics in the oral infection model, both during and after cessation of the RBV treatment. Fecal HEV kinetics were arrested in Phase 1 during the 12-day RBV treatment, similar to the vehicle group (Supplemental Figure S4). On day 30 p.i., which was 18 days after discontinuation of the RBV treatment, 75% of the rats initially treated with RBV still had fecal viral RNA levels below or at the lower limit of quantification (Supplemental Figure S4), whereas levels had increased to detectable levels in 25% of the RBV-treated rats, albeit markedly lower than those in the vehicle-treated rats.

To determine the optimal dose of ribavirin for inhibiting rHEV replication in I.V.-infected athymic nude rats, different doses of RBV (30, 60, or 90 mg/kg/day, once-daily, oral gavage) were evaluated in these rats (*n*=5, each group). Vehicle-treated, infected rats served as controls. Treatment started on the day of infection (2 hours prior to infection) and continued for twelve consecutive days, except for the 90 mg/kg dose for which treatment was discontinued on day 10 p.i. (Figure 5A) due to toxicity of the drug, which included signs of anemia (pale appearance with decreased body temperature), decreased food and/or water intake, and weight loss (Supplemental Figure S5E). No obvious differences in viral RNA kinetics in the feces were found between the vehicle-treated rats and rats treated with the lowest dose (30 mg/kg) of RBV (Figure 5B-C, Supplemental Figure S5A-B, Supplemental Table S3). In both groups, viral RNA kinetics in feces passed through Phase 1 and 2, resulting in similar fecal (Figure 5F) and liver (Figure 5G) viral RNA levels on day 12 p.i.. By contrast, viral RNA kinetics in the feces of rats treated with 60 or 90 mg/kg of RBV were halted in Phase 1 (LLOQ phase, Figure 5D-E, Supplemental Figure S5C-D), resulting in significant reductions (1.87 log_10_ and 1.90 log_10_, respectively; *p*-values ≤ 0.05) in the fecal viral RNA levels on day 12 p.i. as compared to those in the vehicle-treated rats (Figure 5F). This corresponded with a substantially lower viral RNA load (~3 log_10_) in the liver of rats treated with the medium and high dose of RBV (Figure 5G). Due to the adverse effects of the 90 mg/kg dose, the optimal daily dose of RBV for inhibiting rHEV replication in the I.V. infection model is set at 60 mg/kg, although 80% of the rats treated with this dose had detectable HEV RNA in the liver at the end of the 12-day treatment period (Figure 5F, G).

**FIGURE 5.**
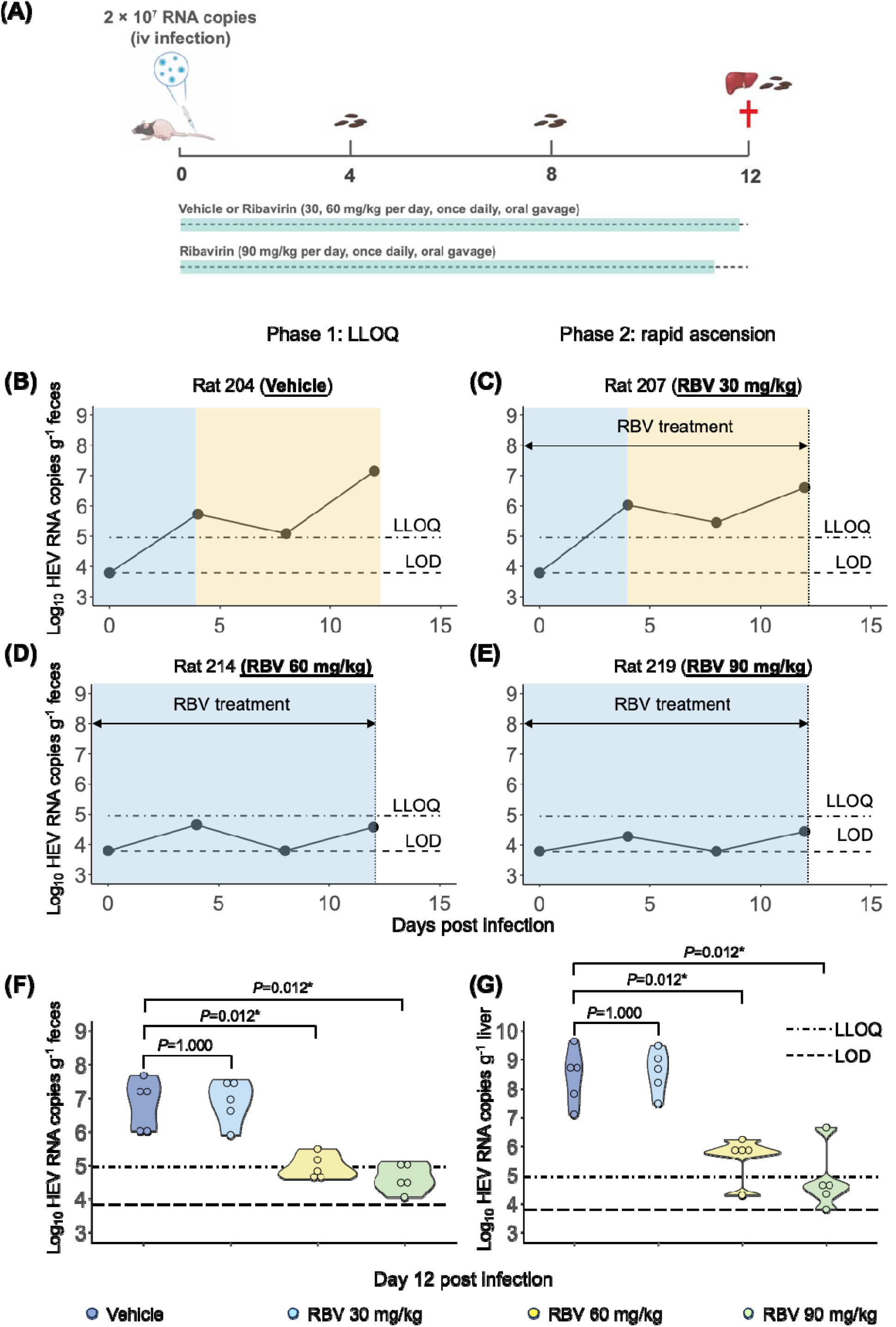
Exploring the optimal dosage of ribavirin in inhibiting rHEV in the I.V. infection model. (A) Schematic representation of evaluating the antiviral effect of RBV at different dosages in the rHEV I.V. infection model. Rat received RBV treatment with 30, 60, and 90 mg/kg, once daily by oral gavage for 12 days (*n*=5 in each group, 11 days of the 90 mg/kg treatment group). The initial dose was administered 2 hours prior to the rHEV infection. (B-E) rHEV kinetic patterns in feces of four representative rats that were treated with (rats 207, 214, and 219) and without RBV (rat 204). Blue shading area and yellow shading area represent Phase 1 and Phase 2, respectively. (F) The viral RNA levels in the feces of individual rats (indicated by solid circles) treated with different dosages of RBV (30, 60, or 90 mg/kg/day) or vehicle on day 12 p.i. are represented in a violin plot. (G) A violine plot illustrating the viral RNA levels in livers treated with various dosages of RBV (30, 60, or 90 mg/kg/day) or vehicle on day 12 p.i.. Individual rats are indicated by solid circles. The LLOQ is depicted by a dotted-dashed line, while the LOD is illustrated using a dashed line. The Kruskal-Wallis Test was employed to evaluate significant differences in HEV RNA levels. *p*-values <0.05 were regarded as statistically significant.

## DISCUSSION

HEV infection is a significant global health burden, especially for high-risk populations, and is primarily transmitted through the fecal-oral route. In this paper, we explored the detailed kinetics of rHEV infection in two rHEV infection models, where rats were either intravenously or orally infected to mimic TT-HEV or HEV transmission via the fecal-oral route, respectively.

The kinetics pattern of rHEV RNA in the feces from initial infection to steady state in the oral infection model revealed a multiphasic rHEV expansion pattern. Following undetectable viral RNA levels in the LLOQ phase (Phase 1), rHEV replicated massively during Phase 2. This was subsequently followed by a relatively steady replication in Phase 3, and a short-term Phase 4 with a sharp increase in the replication rate. Finally, rHEV replication reached a steady phase (Phase 5) until the end of the study. Besides detecting viral RNA in the feces, rHEV RNA levels in the liver during the different phases are consistently ~2 log higher than in the other tissues studied (intestine and spleen). This again reinforces that the liver is the primary site of HEV replication [27], and may serve as the main contributor to the multiphasic replication profile of rHEV due to multiple replication rounds in this tissue. Interestingly, we previously identified a multiphasic kinetic pattern in serum for acute hepatitis B virus (HBV) in a humanized-liver mouse model [28]. In this model, HBV infection and predicted production cycles of progeny virus are exclusively confined to the humanized liver [29], further substantiating that the liver could be solely responsible for the observed multiphasic viral replication kinetics in both infection models of rHEV.

Although primarily hepatotropic, HEV has been associated with various extrahepatic manifestations [27, 30-32]. Moreover, several reports indicate that HEV can replicate in these tissues, including the gastrointestinal tract, kidneys, central nervous system, and placenta [33-35]. By employing human intestinal enteroids, it was recently proposed that HEV infects proliferative cells within the intestinal epithelium [36]. Therefore, we cannot rule out that the multiphasic rHEV kinetic pattern in feces could result from a productive infection in (some of) these organs, followed by viral dissemination from these organs into the feces. Alternatively, the multiphasic kinetic pattern could be the result of a combined contribution of rHEV replication in the hepatic and extrahepatic tissues, with the first 3 phases reflecting rHEV replication in the liver (as seen in HBV-infected mice with humanized livers [28]), while the second increase in viral RNA levels and the subsequent plateau (i.e., Phases 4 and 5) could result from extrahepatic rHEV replication. Since highly sensitive and specific antibodies for the detection of rHEV antigens in tissues are currently lacking, further exploration of other techniques (such as RNAscope) are needed to unequivocally demonstrate that rHEV can productively replicate in tissues other than the liver and to identify the cell types that are implicated in rHEV replication.

The I.V. infection model provides a unique opportunity to examine how the early phases in the rHEV kinetics may be affected compared to those in the oral infection model. While both models display the same initial three phases in the rHEV kinetics, the oral infection model showed an LLOQ phase (Phase 1) that lasted approximately 2 weeks longer than the corresponding phase in the I.V. model. Phases 2 and 3, on the other hand, were similar to those in the oral infection model in terms of duration, viral load, and increase in the slope. The longer Phase 1 is likely due to the initial interaction between the host and rHEV in the gut before the virus reaches the liver via the portal vein in the oral infection model. By contrast, in the I.V. infection model, the virus directly reaches the liver via the blood circulation [30]. Despite the longer Phase 1, the similarity in the initial phases between the two rHEV infection models suggests that the viral replication dynamics are alike once virus replication in the liver has commenced, regardless of the route of infection. Interestingly, the incubation time of rHEV in the oral infection model aligns with the minimal incubation period of 2-weeks reported for HEV GT1 [37, 38], suggesting that oral rHEV infections in rats closely mimic fecal-oral HEV infections in humans.

Because most cases of HEV viremic blood donors are asymptomatic, screening for HEV RNA in blood has been recommended as a viable option to control TT-HEV transmission [39]. Since serologic assays can be unreliable in immunocompromised patients, detection of HEV RNA in blood or stool represents an important (golden) diagnostic method to define an HEV infection [6]. With nucleic acid amplification testing, HEV is detectable in fecal and blood samples at 2 and ~4-5 week after infection, respectively [40]. Our results demonstrate that fecal rHEV RNA levels accurately reflect those in other tissues such as liver, intestine, spleen, and, to a lesser extent those in the serum. Notably, fecal rHEV RNA levels were detected earlier than those in the serum, providing additional evidence that measuring rHEV RNA levels in stool samples could be a cost-effective way for the early detection of an HEV infection in humans and for assessing the rHEV RNA levels in essential tissue compartments. Ribavirin is the only available off-label antiviral treatment, recommended specifically for transplant recipients. Despite the reported efficacy of RBV in patients and humanized mouse models [26, 41, 42], none of the clinical trials conducted thus far have evaluated the most optimal RBV regime [43], resulting in differences in treatment duration and RBV dosage across studies. By studying the viral RNA kinetics, we evaluated the efficacy of different doses of RBV treatment in inhibiting rHEV replication. We here identified 60 mg/kg/day RBV as the optimal dose in the I.V. infection model, corresponding to a median human dose of 583 mg/day of RBV (as determined by a simple dose conversion rule [44, 45]), which is reminiscent of the median dose of 600 (range: 200-1200) mg/day given to patients with a chronic HEV infection after solid organ transplantation [26]. During the 12-day treatment period of the rHEV-infected rats, RBV at a dose 60 mg/kg/day suppressed viral RNA levels in the feces to (nearly) undetectable levels in both the oral infection and the I.V. infection model. In the oral infection model, however, 2 out of 8 rats became positive for rHEV RNA in the feces 18 days after cessation of the 12-day treatment course with 60 mg/kg/day ribavirin. Moreover, this dose did not effectively suppress rHEV levels in the liver of most I.V.-infected rats at the end of the 12-day treatment period. Clearly, this RBV regimen was unable to lower the viral RNA levels to low enough levels to prevent relapse of rHEV, resulting in rapid viral replication (Phase 2) upon treatment discontinuation in both rat HEV infection models. Although the higher (i.e., 90 mg/kg/day) dose was more effective in suppressing rHEV levels in the feces and liver in the I.V. infection model, the rats also experienced serious side effects necessitating early termination of the RBV treatment. These findings suggest that the duration of the RBV treatment may be extended, either using the 60 mg/kg/d-dose or an intermediate dose (i.e., between 60 and 90 mg/kg/day), to potentially avert viral relapse. Alternatively, combining ribavirin with an anti-HEV molecule that operates additively/synergistically to ribavirin may prove to be more effective, as was recently shown [46]. Combination therapy will enable dose reductions, resulting in fewer/no side effects, and reduce the risk of developing drug-induced mutations. More research is needed to evaluate the effectiveness of RBV administration after achieving steady state rHEV RNA levels in the (chronic) rHEV infection model in rats, as was previously done in the (acute) humanized mouse model for HEV [42].

In conclusion, the detailed characterization of the rHEV RNA kinetics reveals important insights regarding the course of the rHEV infection in the rat model. The fecal RNA kinetics showed good concordance with those in various tissue compartments, suggesting that the viral RNA levels in the feces can be good indicators of the status of the rHEV infection. In addition, our study of the rHEV RNA kinetics in different tissue compartments revealed that rHEV appears to replicate first in the liver and then disseminates/replicates in the spleen and intestines, indicating that the liver acts as a reservoir for rHEV. This information will serve as an important foundation for further elucidating the role of the liver as well as the extrahepatic tissues in rHEV replication, which will enable future mechanistic studies and theoretical modelling necessary to elucidate the *in vivo* dynamics of rHEV infection, either or not during treatment.

## Abbreviations

HEV: hepatitis E virus
HBV: hepatitis B virus
rHEV: rat HEV
I.V: intravenous
RBV: ribavirin
TT-HEV: transfusion-transmitted HEV
LLOQ: lower limit of quantification
LOD: limit of detection

## Acknowledgments

We thank Stijn Hendrickx for excellent technical assistance with the animal work and the staff at the Rega animal facility at KU Leuven for technical assistance with the animal experiments.

