## Supplemental Files for "Rat hepatitis E virus infection has multiphasic viral replication kinetics *in vivo*"

**Contents**

**Supplemental Table S1.** Characteristics of rHEV kinetics in the 60-day oral infection model.

**Supplemental Table S2.** Characteristics of rHEV kinetics in the 26-day I.V. infection model.

**Supplemental Table S3.** Characteristics of rHEV kinetics in the 30-day oral infection model upon RBV treatment.

**Supplemental Table S4.** Characteristics of rHEV kinetics in the 12-day I.V. infection model upon RBV treatment.

**Supplemental Figure S1.** rHEV viral kinetic pattern in feces of all rats upon oral infection.

**Supplemental Figure S2.** The linear relationship between median rHEV RNA levels in feces and liver (A), feces and intestine (B), feces and spleen (C), and feces and serum (D) on the studied days.

**Supplemental Figure S3.** rHEV viral kinetics pattern in feces of rat no. 5 from the I.V. infection model (26 days).

**Supplemental Figure S4.** Assessment of the optimal dosage of ribavirin in inhibiting rHEV infection in the oral infection model.

**Supplemental Figure S5.** rHEV kinetic patterns in feces of all I.V.-infected rats.

**Supplemental Table S1.** Characteristics of rHEV kinetics in the 60-day oral infection model

| Rat No.^a^ | Phase 1  (LLOQ) | Phase 2  (Ascension)^b^ | | | Phase 3  (Plateau)^c^ | | Phase 4  (Ascension)^b^ | | | Phase 5  (Plateau)^b^ | |
| --- | --- | --- | --- | --- | --- | --- | --- | --- | --- | --- | --- |
|  | Duration  (days) | Viral load^d^  (log_10_ copies g^-1^ feces) | Slope  (log/day) | Duration  (days) | Viral load^d^  (log_10_ copies g^-1^ feces) | Duration  (days) | Viral load^d^  (log_10_ copies g^-1^ feces) | Slope  (log/day) | Duration  (days) | Viral load^c^  (log_10_ copies g^-1^ feces) | Duration  (days) |
| 17 | 15 | 5.53 | 0.17 | 15 | 8.12 | 5 | 8.11 | 0.32 | 3 | 9.07^e^ | 22 |
| 18 | 15 | 6.34 | 0.13 | 10 | 7.61 | 10 | 8.08 | 0.41 | 3 | 9.29^e^ | 22 |
| 19 | 20 | 5.59 | 0.32 | 5 | 7.17 | 10 | 7.43 | 0.55 | 3 | 9.09 | 22 |
| 20 | 20 | 6.01 | 0.15 | 10 | 7.51 | 5 | 7.68 | 0.23 | 3 | 8.38 | 22 |
| Average  (SD) | 17.50  (2.89) | 5.87  (0.38) | 0.19  (0.087) | 10.00  (4.08) | 7.60  (0.39) | 7.50  (2.89) | 7.83  (0.33) | 0.38  (0.14) | 3.00  (0.00) | 8.74  (0.50) | 22.00  (0.00) |
| Median  [Min-Max] | 17.50  [15-20] | 5.8  [5.53-6.34] | 0.16  [0.13-0.32] | 10  [5-10] | 7.56  [7.17-8.12] | 7.5  [5-10] | 7.88  [7.43-8.11] | 0.37  [0.23-0.55] | 3  [3-3] | 9.08  [8.38-9.29] | 22  [22-22] |
| ^a^Only rats that were followed up until day 60 p.i. are depicted in this table.  ^b^Ascension means slope is significantly different than 0 (*p*-value < 0.05).  ^c^Plateau means slope is not significantly different than 0 (*p*-value > 0.05).  ^d^Viral load is the first viral measurement for each phase.  ^e^Extremely slow decline phase. | | | | | | | | | | | |

**Supplemental Table S2.** Characteristics of rHEV kinetics in the 26-day I.V. infection model

| Rat No.^a^ | Phase 1  (LLOQ) | Phase 2  (Ascension)^b^ | | | Phase 3  (Plateau)^c^ | |
| --- | --- | --- | --- | --- | --- | --- |
|  | Duration  (days) | Viral load^d^  (log_10_ copies g^-1^ feces) | Slope  (log/day) | Duration  (days) | Viral load^d^  (log_10_ copies g^-1^ feces) | Duration  (days) |
| 1 | 6 | 5.64 | 0.17 | 12 | 7.53^e^ | 7^f^ |
| 2 | 2 | 5.24 | 0.19 | 16 | 8.43 | 7 |
| 3 | 2 | 6.18 | 0.12 | 16 | 7.42 | 7 |
| 4 | 2 | 6.13 | 0.15 | 16 | 8.58 | 7 |
| 6 | 2 | 6.47 | 0.18 | 11 | 8.42^e^ | 12^f^ |
| 7 | 8 | 4.82 | 0.24 | 10 | 7.29 | 8 |
| 8 | 6 | 5.70 | 0.18 | 12 | 7.63 | 7 |
| 9 | 4 | 5.01 | 0.17 | 16 | 7.78 | 5 |
| 10 | 4 | 5.58 | 0.16 | 11 | 7.24 | 10 |
| Average  (SD) | 4.00  (2.24) | 5.64  (0.55) | 0.17  (0.03) | 13.33  (2.60) | 7.81  (0.53) | 7.29  (1.50) |
| Median  [Min-Max] | 4  [2-8] | 5.64  [4.82-6.47] | 0.17  [0.12-0.24] | 12  [10-16] | 7.63  [7.24-8.58] | 7  [5-12] |
| ^a^In the 26-day I.V. infection experiment, rat No. 5 were removed due to its distinct kinetic pattern (staircase) (Supplemental Figure S3).  ^b^Ascension means slope is significantly different than 0 (*p*-value < 0.05).  ^c^Plateau means slope is not significantly different than 0 (*p*-value > 0.05).  ^d^Viral load is the first viral measurement for each phase.  ^e^The phase 3 for rat 1 and rat 6 excluded the last data point (day 26). When the last data point was included, the slope of linear regression model is significantly different than 0 (*p*-value < 0.05) (as shown in Figure 4).  ^f^ The duration of phase 4 for rat 1 and rat 6 is calculated by excluding the last data point. | | | | | | |

**Supplemental Table S3.** Characteristics of rHEV kinetics in the 30-day oral infection model upon RBV treatment

| Rat No.^a^ | Phase 1  (LLOQ) | Phase 2  (Ascension)^b^ | | | Phase 3  (Plateau)^c^ | | Phase 4  (Ascension)^b^ | | | Phase 5  (Plateau)^c^ | |
| --- | --- | --- | --- | --- | --- | --- | --- | --- | --- | --- | --- |
|  | Duration  (days) | Viral load^d^  (log_10_ copies g^-1^ feces) | Slope  (log/day) | Duration  (days) | Viral load^d^  (log_10_ copies g^-1^ feces) | Duration  (days) | Viral load^d^  (log_10_ copies g^-1^ feces) | Slope  (log/day) | Duration  (days) | Viral load^d^  (log_10_ copies g^-1^ feces) | Duration  (days) |
| 269 | 16 | 5.77 | 0.22 | 7 | 7.28 | 7 | NA | NA | NA | NA | NA |
| 270 | 12 | 5.26 | 0.20 | 11 | 7.41 | 7 | NA | NA | NA | NA | NA |
| 271 | 16 | 6.32 | 0.11 | 7 | 6.32 | 7 | NA | NA | NA | NA | NA |
| 272 | 18 | 6.51 | 0.16 | 5 | 7.33 | 7 | NA | NA | NA | NA | NA |
| 9 | 19 | 5.37 | 0.15 | 11 | NA | NA | NA | NA | NA | NA | NA |
| 10 | 12 | 5.07 | 0.24 | 11 | 7.69 | 7 | NA | NA | NA | NA | NA |
| 11 | 16 | 6.24 | 0.10 | 11 | 7.39 | 3 | NA | NA | NA | NA | NA |
| 12 | 16 | 5.45 | 0.29 | 7 | 7.49 | 7 | NA | NA | NA | NA | NA |
| Average  (SD) | 15.63  (2.50) | 5.75  (0.55) | 0.18  (0.066) | 8.75  (2.49) | 7.27  (0.44) | 6.43  (1.51) | NA | NA | NA | NA | NA |
| Median  [Min-Max] | 16  [12-19] | 5.61  [5.07-6.51] | 0.18  [0.10-0.29] | 9  [5-11] | 7.39  [6.32-7.69] | 7  [3-7] | NA | NA | NA | NA | NA |
| ^a^The HEV viral RNA levels in feces for all rats treated with 60 mg/kg RBV (Supplemental Figure S4C) are below LLOQ, thus are not depicted in the table.  ^b^Ascension means slope is significantly different than 0 (*p*-value <0.05).  ^c^Plateau means slope is not significantly different than 0 (*p*-value >0.05).  ^d^Viral load is the first viral measurement for each phase.  Abbreviation: NA, not available. | | | | | | | | | | | |

**Supplemental Table S4.** Characteristics of rHEV kinetics in the 12-day I.V. infection model upon RBV treatment

| Rat No.^a^ | Phase 1  (LLOQ) | Phase 2  (Ascension)^b^ | | | Phase 3  (Plateau)^c^ | | Phase 4  (Ascension)^b^ | | | Phase 5  (Plateau)^c^ | |
| --- | --- | --- | --- | --- | --- | --- | --- | --- | --- | --- | --- |
|  | Duration  (days) | Viral load^d^  (log_10_ copies g^-1^ feces) | Slope  (log/day) | Duration  (days) | Viral load^d^  (log_10_ copies g^-1^ feces) | Duration  (days) | Viral load^d^  (log_10_ copies g^-1^ feces) | Slope  (log/day) | Duration  (days) | Viral load^d^  (log_10_ copies g^-1^ feces) | Duration  (days) |
| 201 | 4 | 6.55 | 0.14 | 8 | NA | NA | NA | NA | NA | NA | NA |
| 202 | 4 | 5.40 | 0.23 | 8 | NA | NA | NA | NA | NA | NA | NA |
| 203 | 8 | NA | NA | NA | NA | NA | NA | NA | NA | NA | NA |
| 204 | 4 | 5.73 | 0.18 | 8 | NA | NA | NA | NA | NA | NA | NA |
| 205 | 8 | NA | NA | NA | NA | NA | NA | NA | NA | NA | NA |
| 206 | 4 | 6.40 | 0.07 | 8 | NA | NA | NA | NA | NA | NA | NA |
| 207 | 4 | 6.03 | 0.07 | 8 | NA | NA | NA | NA | NA | NA | NA |
| 208 | 12 | NA | NA | NA | NA | NA | NA | NA | NA | NA | NA |
| 209 | 4 | 6.49 | 0.11 | 8 | NA | NA | NA | NA | NA | NA | NA |
| 210 | 4 | 6.43 | 0.14 | 8 | NA | NA | NA | NA | NA | NA | NA |
| Average  (SD) | 5.60  (2.80) | 6.15  (0.44) | 0.13  (0.06) | 8  (0) | NA | NA | NA | NA | NA | NA | NA |
| Median  [Min-Max] | 4  [4-12] | 6.40  [5.40-6.55] | 0.14  [0.07-0.23] | 8  [8-8] | NA | NA | NA | NA | NA | NA | NA |
| ^a^In the 12-day I.V infection experiment, rat No. 206-210 were treated with ribavirin (30 mg/kg) (Supplemental Figure S6). Two sample Mann-Whitney *U* test shows that there is no significantly difference (*p*-value >0.05) between this group and the vehicle group for each phase (in terms of duration, initial viral load, and slope for phase 1 and phase 2). Only rats that were followed up until day 12 p.i. are depicted in the table.  In the 12-day I.V infection experiment, all rats (five rats) treated with ribavirin (60 mg/kg) were in phase 1 for 12 days (Supplemental Figure S7). Rats 211 and 212 had significant levels of viral load only on day 12. Rat 213 had a significant level of viral load only on day 4 and then went down to LLOQ for the remaining 8 days.  ^b^Ascension means slope is significantly different than 0 (*p*-value <0.05).  ^c^Plateau means slope is not significantly different than 0 (*p*-value >0.05).  ^d^Viral load is the first viral measurement for each phase.  Abbreviation: NA, not available. | | | | | | | | | | | |

### Supplemental Figure S1

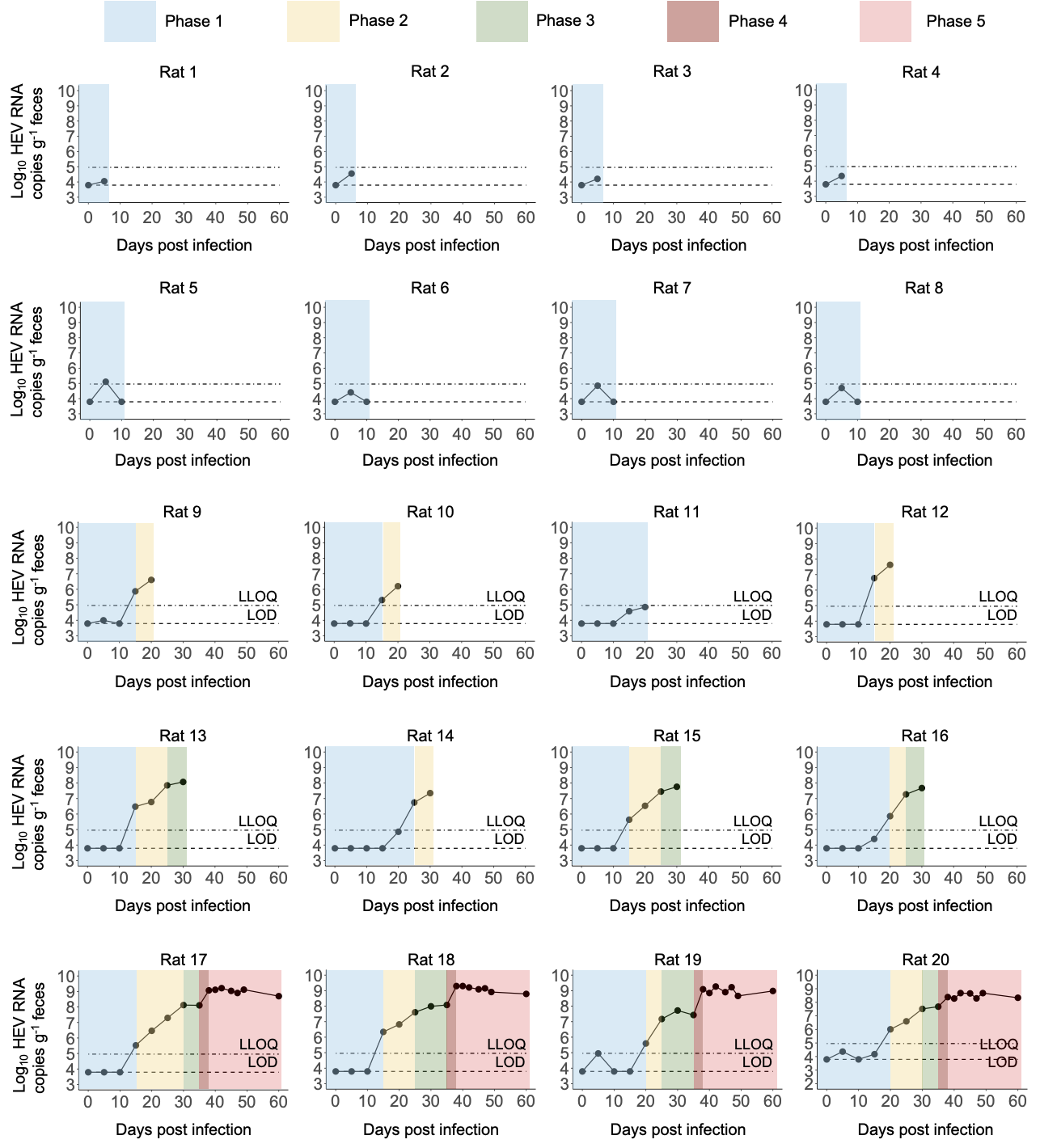

**Supplemental Figure S1.** rHEV viral kinetic pattern in feces of all rats (*n*=20) upon oral infection. LLOQ and LOD indicate lower limit of quantification and limit of detection, respectively.

**Supplemental Figure S2**

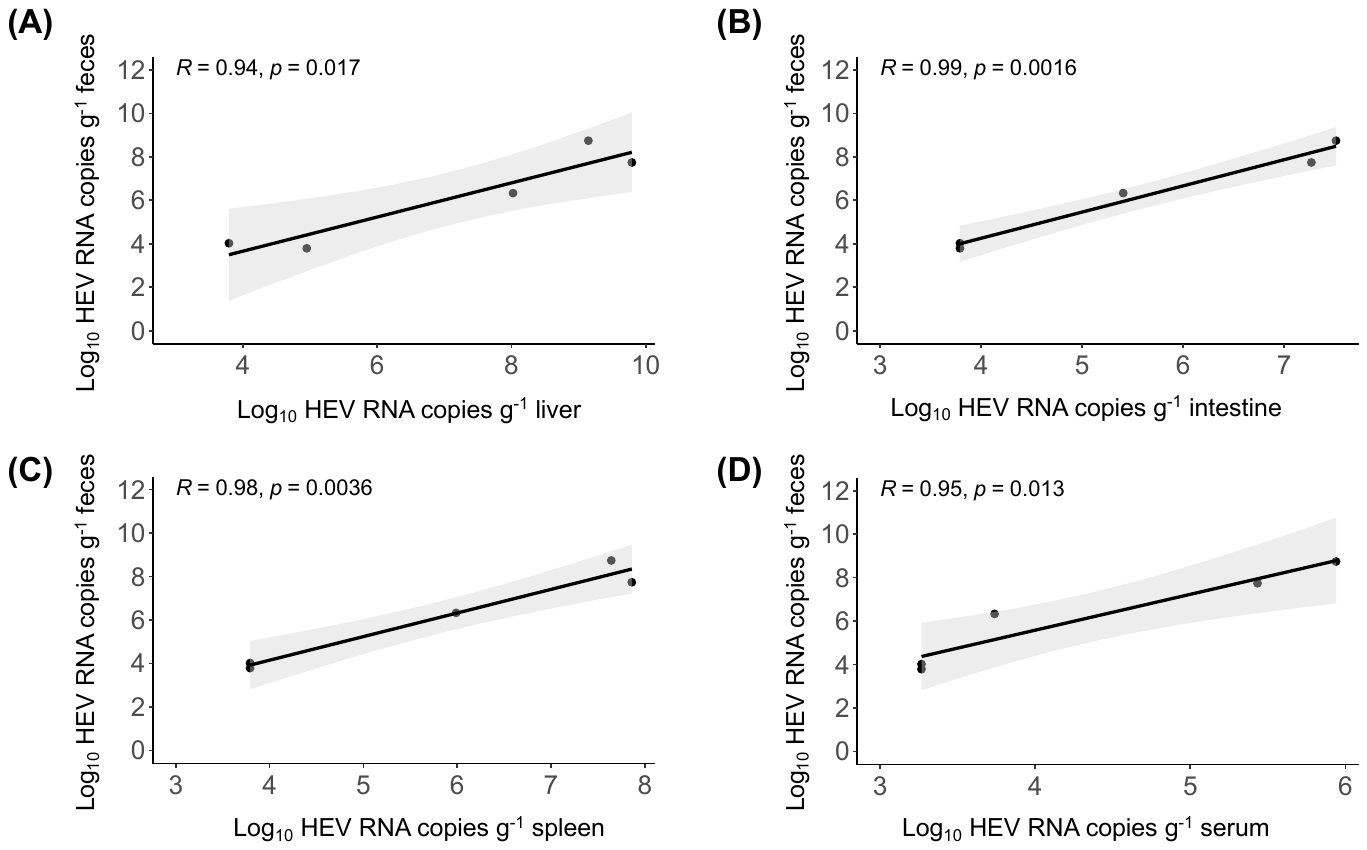

**Supplemental Figure S2.** The linear relationship between median rHEV RNA levels in feces and liver (A), feces and intestine (B), feces and spleen (C), and feces and serum (D) on the studied days. Data points below LOD were set to 4 log_10_, which is the assay threshold. Each dot represents a pair of median rHEV RNA levels at different days post oral infection, *R* represents the Pearson correlation coefficient, and *p* is the corresponding 2-sided *p*-value.

**Supplemental Figure S3**

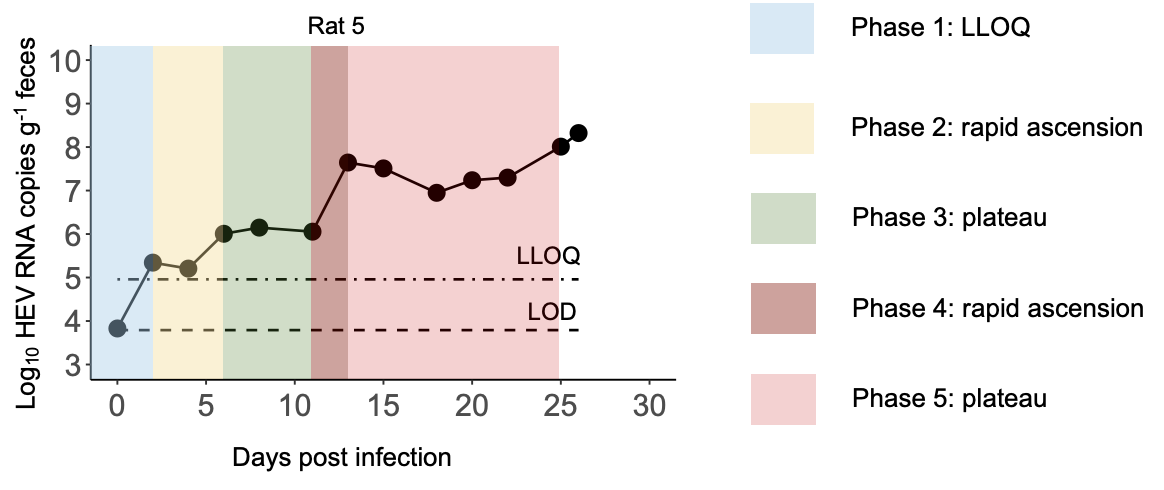

**Supplemental Figure S3.** rHEV viral kinetics pattern in feces of rat no. 5 from the I.V. infection model (26 days). Compared to Figure 4, only two transit phases were observed (Phase 3: plateau; and Phase 4: rapid ascension) for rat no. 5. LLOQ and LOD indicate lower limit of quantification and limit of detection, respectively.

**Supplemental Figure S4**

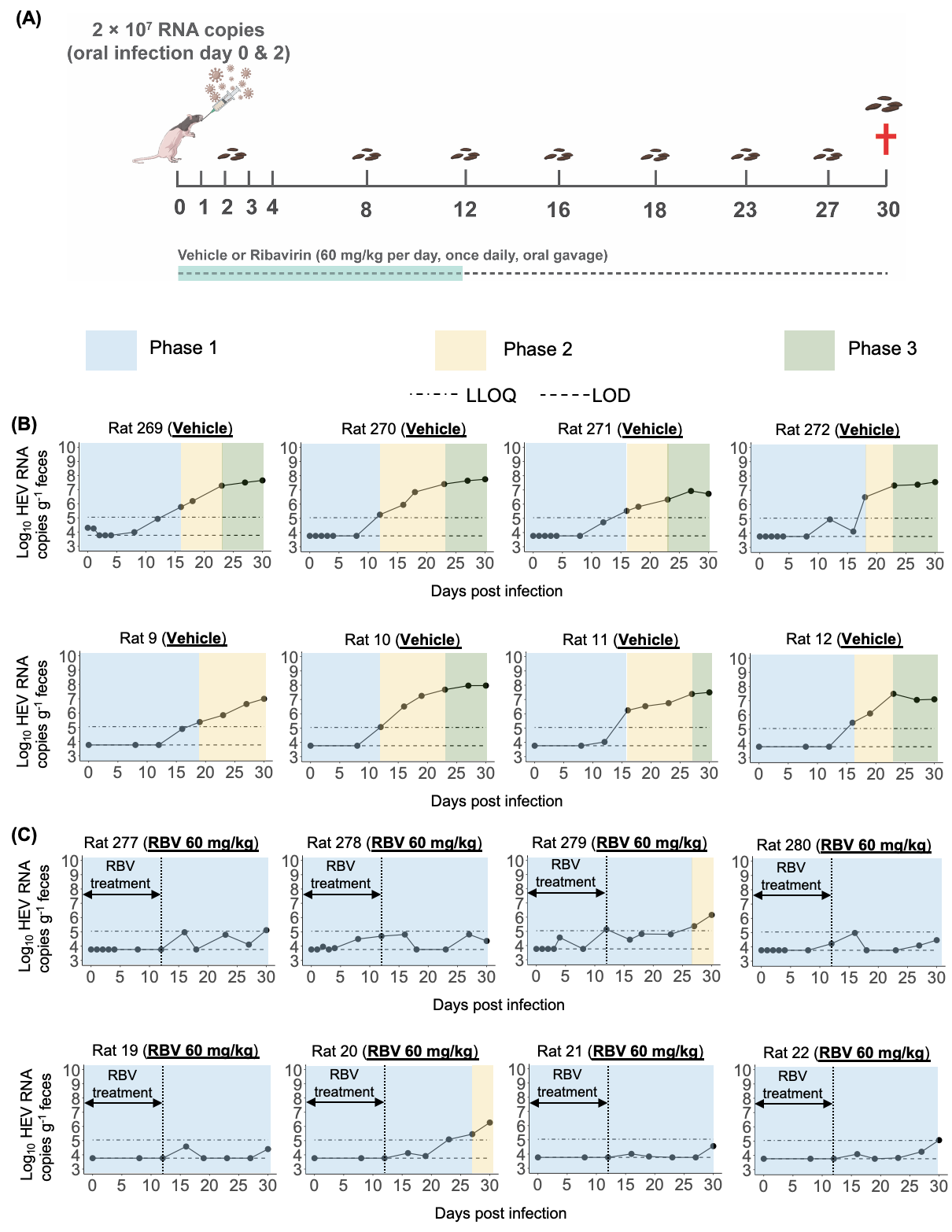

**Supplemental Figure S4.** Assessment of the optimal dosage of ribavirin in inhibiting rHEV infection in the oral infection model. (A) Diagram illustrating the evaluation of the antiviral effect of RBV 60 mg/kg in the oral infection model. Rats (*n*=16, *n*=8/group) were infected orally with a suspension of feces on day 0 and 2 p.i.. Ribavirin was administered orally to the rats at a dosage of 60 mg/kg once daily for 12 days until day 11 p.i., with the first administration occurring 2 hours prior to infection. (B) rHEV kinetic patterns in the feces of rats that received vehicle treatment. (C) rHEV kinetic patterns in the feces of rats that received 60 mg/kg RBV treatment. Blue shading, yellow shading, and green shading areas represent Phase 1, Phase 2, and Phase 3, respectively. The LLOQ is indicated by a dotted-dashed line, while the LOD is represented by a dashed line.

**Supplemental Figure S5**

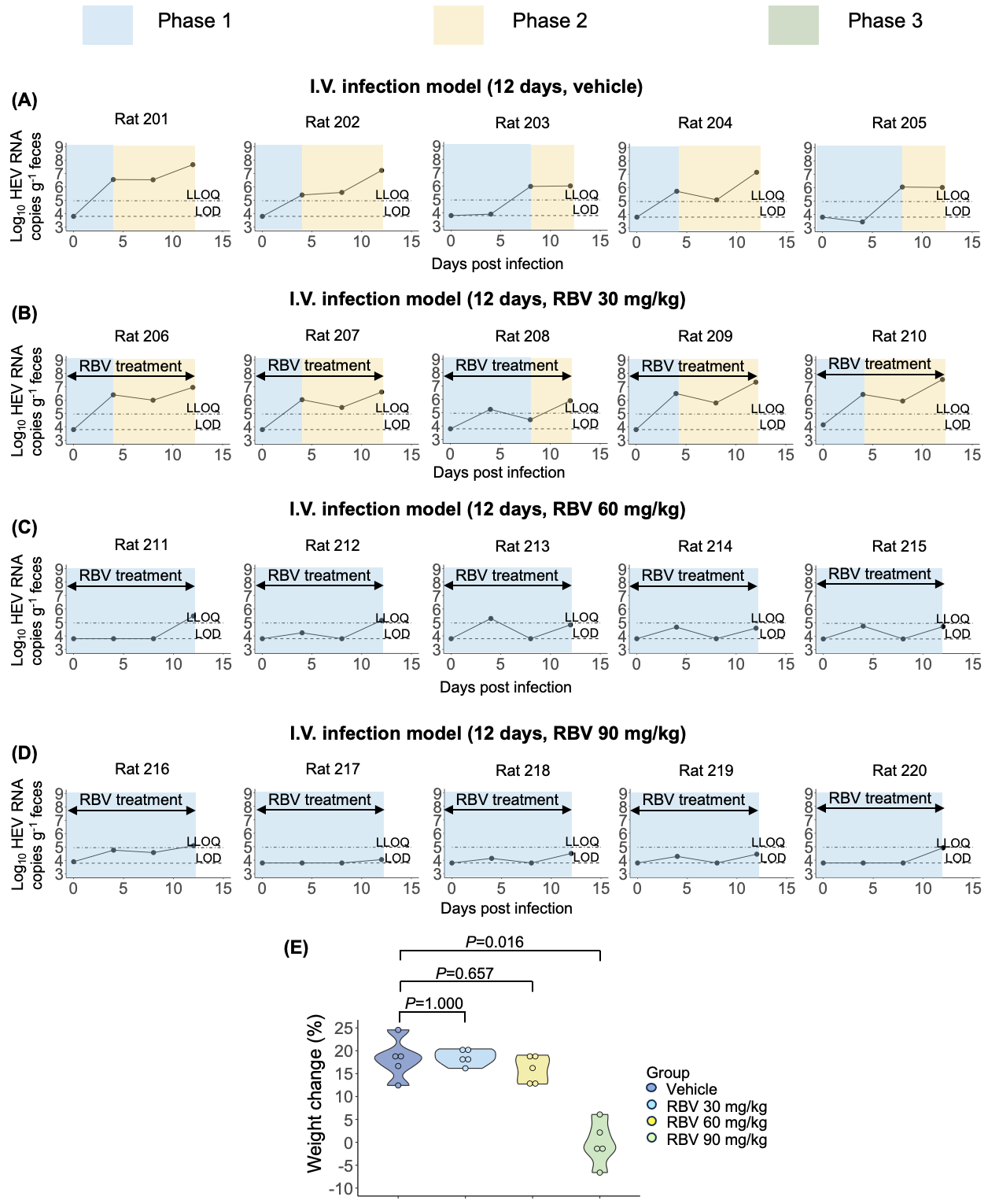

**Supplemental Fig. S5.** rHEV kinetic patterns in feces of all I.V.-infected rats. (A) Rats (*n*=5) that received vehicle treatment. (B) Rats (*n*=5) that received RBV treatment (30 mg/kg). (C) Rats (*n*=5) that received RBV treatment (60 mg/kg). (D) Rats (*n*=5) that received RBV treatment (90 mg/kg). (E) A violin plot illustrating the percentage of weight change for individual rats (indicated by solid circles) on day 12 p.i. relative to their weight on day 0 p.i.. LLOQ (dotted-dashed line) and LOD (dashed line) indicate lower limit of quantification and limit of detection, respectively. The Kruskal-Wallis Test was employed to evaluate significant differences in weight change among the different groups. *p*-values <0.05 were regarded as statistically significant. LLOQ (dotted-dashed line) and LOD (dashed line) indicate lower limit of quantification and limit of detection, respectively.
